# Single-cell analysis of severe COVID-19 patients reveals a monocyte-driven inflammatory storm attenuated by Tocilizumab

**DOI:** 10.1101/2020.04.08.029769

**Authors:** Chuang Guo, Bin Li, Huan Ma, Xiaofang Wang, Pengfei Cai, Qiaoni Yu, Lin Zhu, Liying Jin, Chen Jiang, Jingwen Fang, Qian Liu, Dandan Zong, Wen Zhang, Yichen Lu, Kun Li, Xuyuan Gao, Binqing Fu, Lianxin Liu, Xiaoling Ma, Jianping Weng, Haiming Wei, Tengchuan Jin, Jun Lin, Kun Qu

## Abstract

Despite the current devastation of the COVID-19 pandemic, several recent studies have suggested that the immunosuppressive drug Tocilizumab can powerfully treating inflammatory responses that occur in this disease. Here, by employing single-cell analysis of the immune cell composition of severe-stage COVID-19 patients and these same patients in post Tocilizumab-treatment remission, we have identified a monocyte subpopulation specific to severe disease that contributes to inflammatory storms in COVID-19 patients. Although Tocilizumab treatment attenuated the strong inflammatory immune response, we found that immune cells including plasma B cells and CD8^+^ T cells still exhibited an intense humoral and cell-mediated anti-virus immune response in COVID-19 patients after Tocilizumab treatment. Thus, in addition to providing a rich, very high-resolution data resource about the immune cell distribution at multiple stages of the COVID-19 disease, our work both helps explain Tocilizumab’s powerful therapeutic effects and defines a large number of potential new drug targets related to inflammatory storms.

## Introduction

As of May 1, 2020, the WHO has reported 224,172 deaths out of 3,175,207 confirmed cases for infection by severe acute respiratory syndrome coronavirus 2 (SARS-CoV-2), and these numbers are still growing rapidly^1^. Approximately 14% of patients with COVID-19 experienced severe disease, and 5% were critically ill, among which there was a 49% fatality rate^2^; it has been speculated that this high mortality is related to abnormal immune system activation^3, 4, 5^. Hence, there is an urgent need for researchers to understand how the immune system responds to SARS-CoV-2 viral infection at the severe stage, which may highlight potential effective treatment strategies.

Studies have shown that the inflammatory storm caused by excessive immune responses was strongly associated with mortality in COVID-19^6,7^. Plasma concentrations of a series of inflammatory cytokines, such as granulocyte-macrophage colony-stimulating factor (GM-CSF), interleukin (IL)-6^4^, tumor necrosis factor α (TNF-α), IL-2, 7, 10, and granulocyte colony-stimulating factor (G-CSF)^8^ were increased after SARS-CoV-2 infections. Further investigation demonstrated that peripheral inflammatory monocytes and pathogenic T cells may incite cytokine storm in severe COVID-19 patients^4,6^. Tocilizumab, an immunosuppressive drug that targets IL-6 receptors, has been used to treat severe COVID-19 patients^9,10^, as it is effective for treating severe and even life-threatening cytokine-release syndrome ^11,12^. After receiving Tocilizumab, the body temperature of the patients returned to normal after 24 hours, and Tocilizumab was shown to significantly decrease the concentration of oxygen inhalation by COVID-19 patients by the 5^th^ day of treatment^13^. Despite the apparent efficacy of Tocilizumab for treating severe COVID-19 patients, the lack of single-cell level analyses has prevented any deepening of our understanding about how Tocilizumab impacts the typical COVID-19 induced activation of an inflammatory storm.

In the present study, we profiled the single-cell transcriptomes of 13,239 peripheral blood mononuclear cells (PBMCs) isolated at the severe and remission disease stages of two severe COVID-19 patients treated with Tocilizumab. We identified a severe-stage-specific monocyte subpopulation that clearly contributes to the patients’ inflammatory storms. Comparison between the severe and remission disease stages at the single-cell level revealed that Tocilizumab treatment weakens the excessively activated inflammatory immune response and also showed that immune cells, including plasma B cells and CD8^+^ T cells, still exhibit boosted humoral and cell-mediated anti-virus immune responses in post-treatment COVID-19 patients. Our study thus provides a rich, high-resolution data set about the immune context at multiple stages of COVID-19, and helps to explain how a promising candidate drug both alters immune cell populations and reduces patient mortality.

## Results

### An atlas of peripheral immune cells in severe COVID-19 patients

We obtained 5 peripheral blood samples from 2 severe COVID-19 patients at 3 time points including the severe and remission stages during Tocilizumab treatment (Fig. 1a). Specifically, we collected blood samples at day 1— within 12 hours of Tocilizumab administration—and at day 5 for both patients; note that we also obtained a blood sample from patient P2 on day 7 of Tocilizumab treatment because P2 still had a positive result for a SARS-Cov-2 nucleic acid test of a throat swab specimen on day 5. At day 1, the patients both had a decreased number of lymphocytes compared to healthy reference interval, as well as increased percentages of neutrophils and elevated concentrations of C-reaction protein and increased expression of IL-6 (Supplementary Table 1). Since the clinical symptoms of most of the severe COVID-19 patients, including both patients in this study, were remarkably improved by 5 days of Tocilizumab treatment^13^ (Supplementary Table 1), we defined the blood draws from day 5 as the “remission stage”. For patient P2, we took another blood draw at day 7, when his nucleic acid test turned negative (Fig. 1a). It is worth noting that patient P1 was discharged on day 8, and patient P2 on day10, and these discharges were both at 3 days after a nucleic acid test of a throat swab specimen was negative.

**Figure 1.**
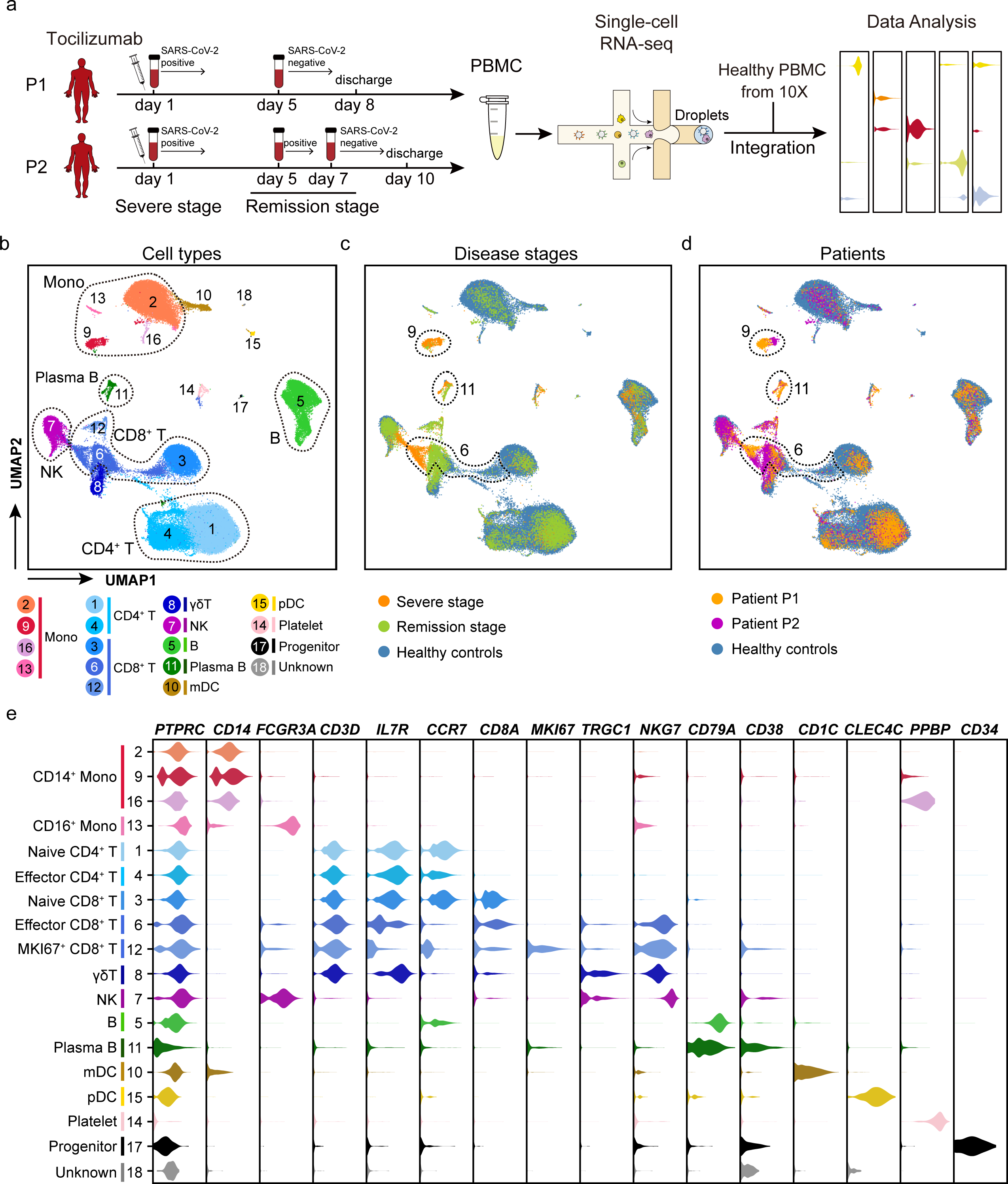
An atlas of peripheral immune cells in severe COVID-19 patients. **a**, Flowchart depicting the overall design of the study. Blood draws from patient P1 were performed at 2 time points (day 1 and day 5), and from P2 at 3 time points (day 1, day 5 and day 7). P1 at day 1 and P2 at day 1 and day 5 were positive for the nucleic acid test of a throat swab specimen. P1 at day 5 and P2 at day 7 were negative for the nucleic acid test of a throat swab specimen. Patients at day 1 were at the severe stage, were in the remission stage at day 5 (P1 and P2); the day 7 blood draw for P2 (still remission stage) was based on a positive nucleic acid test at day 5. Note that samples on day 1 were collected within 12 hours of Tocilizumab treatment. **b-d**, UMAP representations of integrated single-cell transcriptomes of 69,237 PBMCs, with 13,239 cells from our COVID-19 patients and 55,998 were from 10X official website^16^. Cells are color-coded by clusters **(b)**, disease state **(c)**, and sample origin **(d)**. Dotted circles represented cell types with > 5% proportion of PBMCs in **(b)**, and clusters significantly enriched in patients versus controls in **(c, d)**. Mono, monocyte; NK, natural killer cells; mDC, myeloid dendritic cells; pDC, plasmacytoid dendritic cells. **e**, Violin plots of selected marker genes (upper row) for multiple cell subpopulations. The left column presents the cell subtypes as identified based on combinations of marker genes.

We isolated the PBMCs from the COVID-19 patients’ blood samples and subjected them to single-cell mRNA sequencing (scRNA-seq) using the 10X platform (Fig. 1a). After rigorous quality control definition (Supplementary Fig. 1a-d, Supplementary Table 2), low quality cells were filtered; we also removed cell doublets using Scrublet^14^. Correlation of the gene expression for the samples from either patient emphasized the excellent reproducibility between the technical and biological replicates of our dataset (Supplementary Fig. 1e-f). After quality control (QC) and doublet removal, our dataset comprised a total of 13,239 high-quality transcriptomes for single PBMCs.

Due to the similarities between the single-cell transcriptomes of most of the identified cell subsets at the severe and remission disease stages, we initially combined the samples from both patients from day 1 as the “severe stage” and combined the samples from day 5 (and day 7 for P2) as the “remission stage”; note that we also conducted separate analyses for each patient, which yielded similar data trends (Supplementary Fig. 2a, b). In total, the combined analyses of all the single-cell transcriptomes for the COVID-19 patients included 4,344 cells from the severe disease stage and 8,895 were from the remission disease stage.

To investigate heterogeneity among the PBMCs for the COVID-19 patients compared to healthy controls, we applied Seurat^15^ (version 3.1.4) to integrate our COVID-19 single-cell transcriptomes with the published single-cell profiles of healthy PBMCs from the 10X official website^16^, enabling an analysis with a total of 68,190 cells (See **Methods**). We then normalized and clustered the gene expression matrix; this identified 18 unique cell subsets, which were visualized via uniform manifold approximation and projection (UMAP) (Fig. 1b-d). Cell lineages, including monocytes, CD4^+^ and CD8^+^ T, γδT, natural killer (NK), B, plasma B and myeloid dendritic cells (mDC), plasmacytoid dendritic cells (pDC), platelets, and CD34^+^ progenitor cells were identified based on the expression of known marker genes (Fig. 1e). This analysis represents a delineation of the landscape of circulating immune cells for severe COVID-19 patients.

We also used another integration method, Harmony^17^, to help assess the accuracy of the cell clustering results from Seurat^15^ (version 3.1.4) and again visualized the results in UMAP (Supplementary Fig. 3a). We found strong correlations for the identified cell subsets and the detected gene expression patterns between the cell clusters with the two integration methods (Supplementary Fig. 3b, c), supporting the robustness of our cell clustering results.

We next explored the distribution of immune cells from the severe and remission stage COVID-19 patients, as well as in healthy control individuals (Supplementary Fig. 4a). We observed that a number of subpopulations, such as pDCs (cluster 15), mDCs (cluster 10), and most monocytes (clusters 2 and 13) were present in remission stage COVID-19 patients and in healthy controls but not in severe COVID-19 patients (Supplementary Fig. 4b), indicating that Tocilizumab treatment gradually restores a normal distribution of these cell types in circulating blood. Some cell subsets such as NK cells (cluster 7) and CD4^+^ T cells (cluster 1 and 4) were quite heterogeneous between the two COVID-19 patients, so we did not examine these cell types further. These analyses revealed the conspicuous presence of four cell populations that were uniquely present in the COVID-19 patients (albeit to differencing extents in the severe vs. remission disease stages), including a monocyte subpopulation (cluster 9), plasma B cells (cluster 11), effector CD8^+^ T cells (cluster 6), and proliferative MKI67^+^CD8^+^ T cells (cluster 12) patients (Supplementary Fig. 4c). Given our study’s aim of characterizing the COVID-19-specific and Tocilizumab-sensitive immune cell populations of COVID-19 patients, the majority of our subsequent detailed analyses focused on these four cell populations.

### A monocyte subpopulation contributes the inflammatory storm in severe stage COVID-19 patients

Monocytes have been reported to play a vital role in CAR-T induced cytokine-release syndrome^18^ and in SARS-CoV-2 infection triggered inflammatory storms^4^, so we explored the features and functions of the aforementioned monocyte subpopulation that we detected in our single-cell analysis of the two COVID-19 patients. We detected 1,677 monocytes in patients, with 916 from the severe disease stage and 761 from the remission stage; we examined these alongside the data for 9,517 monocytes from health controls. The UMAP plot displayed two main clouds of monocytes that were clearly segregated (Fig. 2a). One monocyte subpopulation (cluster 9) consisted of 98.3% of all monocytes at severe stage, while this ratio was only 12.1% at remission stage and 0% in healthy controls (Fig. 2b), so we initially assessed these severe-stage-specific monocytes.

**Figure 2.**
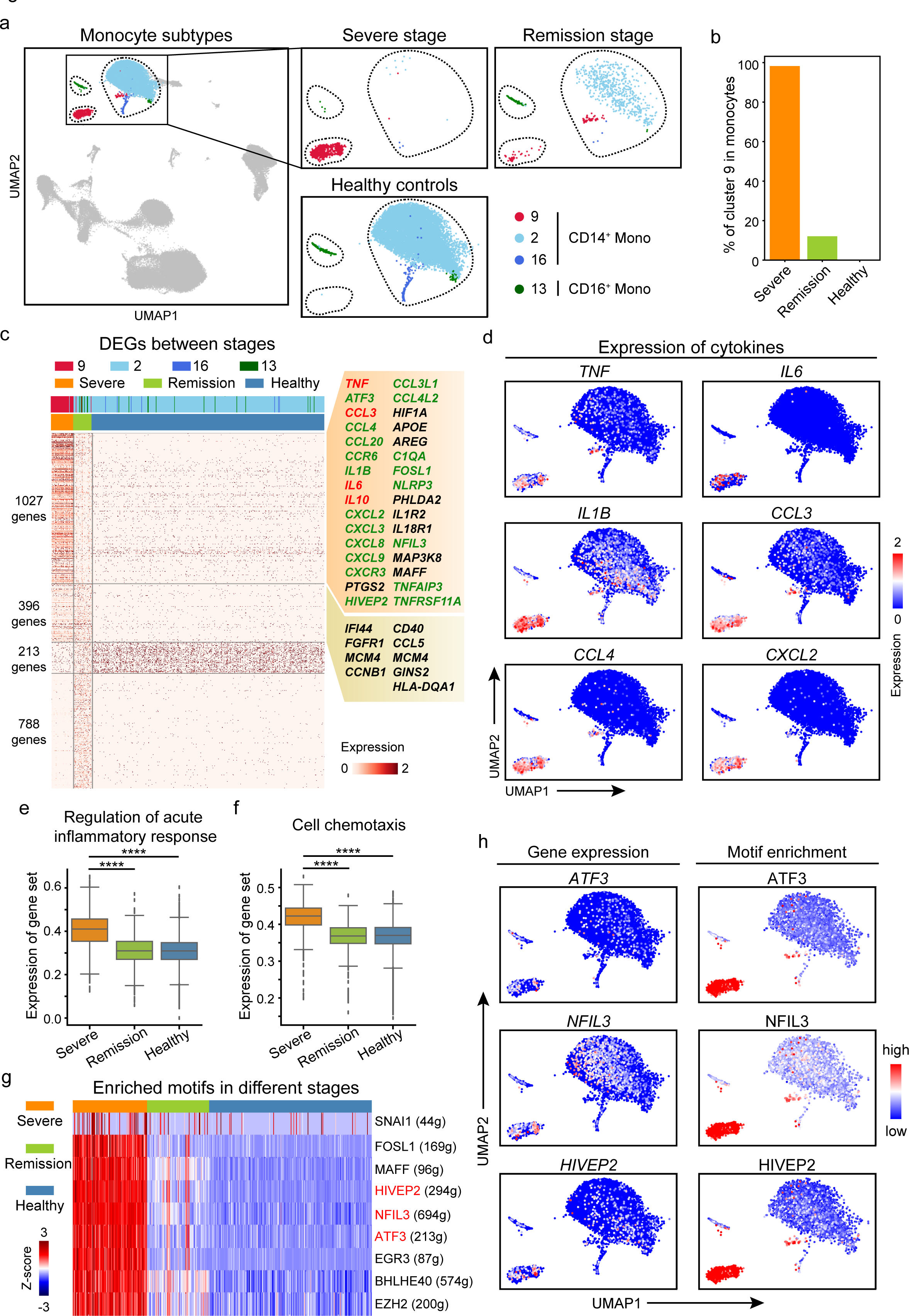
A unique monocyte subpopulation contributes to the inflammatory storm in severe-stage COVID-19 patients. **a**, UMAP plot showing 3 clusters of CD14^+^ monocytes and 1 cluster of CD16^+^ monocyte. Cells are color-coded by clusters. **b**, Bar plot of the proportion of monocytes in cluster 9 at the severe and remission stages, and in healthy control individuals. **c**, Heatmap of differentially expressed genes (DEGs) in monocytes from pairwise comparison between the severe stage patients, remission stage patients, and healthy control individuals. **d**, UMAP plots showing the expression of selected cytokines in all monocyte clusters. **e**,**f**, Box plot of the average expression of genes involved in the signaling pathway “Regulation of acute inflammatory response “and “Cell chemotaxis” in monocytes from the severe and remission stages, and in healthy control individuals. Center line, median; box limits, upper and lower quartiles; whiskers, 1.5x interquartile range; points, outliers; **** represents *P* value < 10^−100^, Wilcoxon rank-sum test. **g**, Heatmap of the area under the curve (AUC) scores of expression regulation by transcription factors (TFs), as estimated using SCENIC. Shown are the top-ranked TFs having the highest difference in expression regulation estimates in monocytes from severe-stage COVID-19 patients. **h**, UMAP plots showing the expression of the *ATF3, NFIL3*, and *HIVEP2* genes in monocytes (top) and the AUC of the estimated regulon activity of the corresponding TFs, predicting the degree of expression regulation of their target genes (bottom).

Transcriptional differences among monocytes subtypes was detected based on a pairwise comparison of the gene expression in the severe and remission stages and in respective comparisons against healthy control individuals. A large number of differentially expressed genes (DEGs) with reported inflammation-related functions were observed in the severe-stage-specific monocytes, including the previously reported cytokine-storm-related genes such as *TNF*^8^, *IL10*^*8*^, *CCL3*^8^, and *IL6*^4^; inflammatory related chemokine genes *CCL4, CCL20, CXCL2, CXCL3, CCL3L1, CCL4L2, CXCL8* and *CXCL9*; and inflammasome activation associated genes *NLRP3* and *IL1B* in the severe-stage-specific monocytes (Fig. 2c, fold change > 2, *P* < 10^−3^, Wilcoxon rank-sum test; Fig. 2d; and Supplementary Table 3). Collectively, the large number of DEGs with reported inflammation-related functions support the idea that the severe-stage-specific monocyte subpopulation we detected in our single-cell COVID-19 patient data may strongly support development of inflammatory responses in severe COVID-19 patients.

A GO analysis indicated enrichment of genes with annotations related to “regulation of acute inflammatory response”, “regulation of leukocyte activation”, “cell chemotaxis” and “cellular response to chemokine” in severe-stage COVID-19 patients compared to remission-stage patients and healthy controls (Fig. 2e, f, *P* < 10^−117^, Wilcoxon rank-sum test; Supplementary Fig. 5; and Supplementary Table 4, 5), suggesting that the inflammatory storm caused by this monocyte subpopulation is suppressed by Tocilizumab treatment.

Next, we explored transcription factors (TFs) in monocytes which may be involved in the promoting of the inflammatory storm. We used SCENIC^19^ and predicted 9 TFs that may regulate genes that were up-regulated in severe-stage-specific monocytes (Fig. 2g). We then constructed a gene regulatory network among the SCENIC predicted TFs and a set of inflammation-relevant genes that were collected from the literatures ^20, 21^. We found that 3 of the SCENIC predicted TFs, namely *ATF3, NFIL3*, and *HIVEP2*, may have the capacity to regulate the detected inflammation-relevant genes (Supplementary Fig. 6). Additionally, we found that the expression of *ATF3, NFIL3*, and *HIVEP2* and their motif enrichment which was predicted by the expressing of their potential target genes were enhanced in the severe-stage-specific monocyte subpopulation (Fig. 2h), further supporting that these 3 TFs may regulate the observed inflammatory storm in monocytes.

Recent studies have shown that over 20% of the severe COVID-19 patients had symptoms of severe septic shock, which affects several organ systems and contributes to liver injury^22^, acute kidney failure^23^, and abnormal heart damage^24^. We therefore checked whether this severe-stage-specific monocyte subpopulation is unique to COVID-19. We downloaded scRNA-seq datasets from patients with sepsis at a mild stage (Int-URO) and patients with sepsis at a severe stage (ICU-SEP), as well as critically ill patients without sepsis (ICU-NoSEP) and healthy controls (Control)^25^. We then integrated these data sets with our COVID-19 patients’ single-cell data using Seurat^15^ (version 3.1.4), which revealed a total of 10 monocyte cell clusters (Supplementary Fig. 7a, b). Interestingly, the cells from the severe stage COVID-19 patients clearly overlapped with only one of the integrated monocyte clusters (cluster VI) (Supplementary Fig. 7c), suggesting that the severe-stage-specific monocyte population might be unique to COVID-19.

### A monocyte-centric cytokine/receptor interaction network in severe-stage COVID-19 patients

Given that monocytes in the severe stage may be involved in the regulation of a variety of immune cell types, we used the accumulated ligand/receptor interaction database^26^ CellPhoneDB (www.cellphonedb.org) to identify alterations of molecular interactions between monocytes and all of the immune cell subsets we identified in our single-cell analysis (Supplementary Table 6). We found 15 cytokine/receptors pairs whose interactions were significantly boosted in severe-stage COVID-19 patients as compared to remission stage patients and healthy controls (Fig. 3a). It is notable that the expression of multiple inflammatory-storm-related cytokines/receptors was significantly increased in severe stage COVID-19 patients (Fig. 3b), which seems plausible that monocytes may have a substantially increased propensity for interaction with other immune cells in blood vessels. Our comparison between severe stage and remission stage patients also suggested obvious attenuation of increased cytokine/receptor interaction activity among the immune cells of remission COVID-19 patients (Fig. 3b). While clearly preliminary, our data support a role for Tocilizumab in reducing monocyte receptor compositions that have been previously implicated in the induction of inflammatory storms.

**Figure 3.**
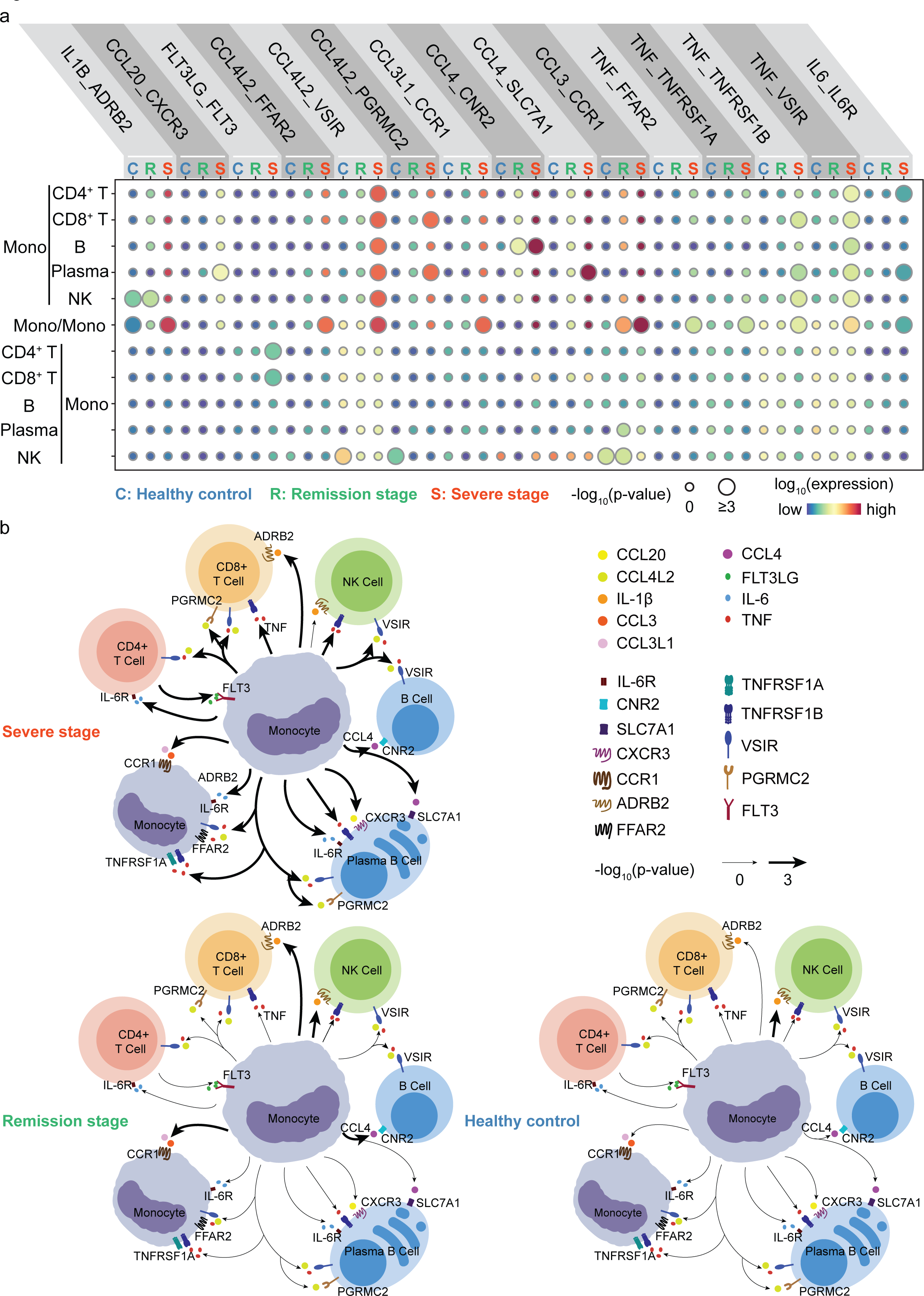
The monocyte-centric molecular interactions of peripheral immune cells in severe-stage COVID-19 patients. **a**, Dot plot of predicted interactions between monocytes and the indicated immune cell types in the severe and remission stages, and in healthy control individuals. *P* values were measured by circle sizes, scale on right (permutation test). The means of the average expression level of interacting molecule 1 in cluster 1 and interacting molecule 2 in cluster 2 are indicated by color. Assays were carried out at the mRNA level, but are extrapolated to protein interactions. **b**, Summary illustration depicting the potential cytokine/receptor interactions between monocytes and other types of peripheral immune cells in the severe and remission stages, and in healthy control individuals. Bolder lines indicate predicted enriched cytokine/receptor interactions between monocytes and other immune cell types.

Consistent with a previous reported that the inflammatory monocytes released IL-6 play an vital role in inciting inflammatory storm in severe COVID-19 patients^4^, we found monocytes were predicted to communicate with CD4^+^ T cells and plasma B cells at severe stage COVID-19 patients through the cytokine/receptor pairs of IL-6/IL-6R. We also detected that the severe-stage-specific monocytes featured elevated expression of other cytokine/receptor pairs that may contribute to a broad spectrum of immune cell communications, such as TNF-α and its receptors, through which monocytes may interact with CD4^+^ T, CD8^+^ T and B cells. Similarly, the severe-stage monocytes had elevated levels of IL-1β and its receptor, suggesting potentially functional interaction of these monocytes with CD8^+^ T cells. Chemokines such as CCL4L2, CCL3, and CCL4 and their respective receptors were also found to be enriched in severe stage monocytes, indicating the potential of targeting these cytokines and/or their receptors as possible drug targets for treating severe-stage COVID-19 patients. Indeed, it is notable that inhibitors targeting some of these cytokine/receptor pairs are currently undergoing anti-COVID-19 clinical trials in multiple places around the world (Supplementary Table 7). Collectively, these findings help illustrate the possible molecular basis of cell-cell interactions at the peripheral blood of COVID-19 patients, leading to a better understanding of the mechanisms of inflammatory storm of the disease.

### Boosted humoral and cell-mediated immunity in severe COVID-19 patients

Studies of avian H7N9 disease have revealed that viral infection can elicit robust, multi-factorial immune responses^27,28^, and a very recent study reported effective immune responses from a non-severe COVID-19 patient^29^. However, it is not clear whether the anti-virus immune responses are affected by Tocilizumab treatment. We assessed the anti-virus immune responses—both humoral and cell-mediated immune responses—of severe-stage COVID-19 patients as compared with both remission stage patients and healthy controls. As expected for un-infected controls, there were hardly any plasma B cells in healthy individuals (Fig. 4a). In contrast, there were many plasma B cells in both the severe and remission stage COVID-19 patients (Fig. 4a, b), suggesting that SARS-CoV-2 infection may elicit the anti-virus humoral immune responses, which are not affected by the Tocilizumab treatment.

**Figure 4.**
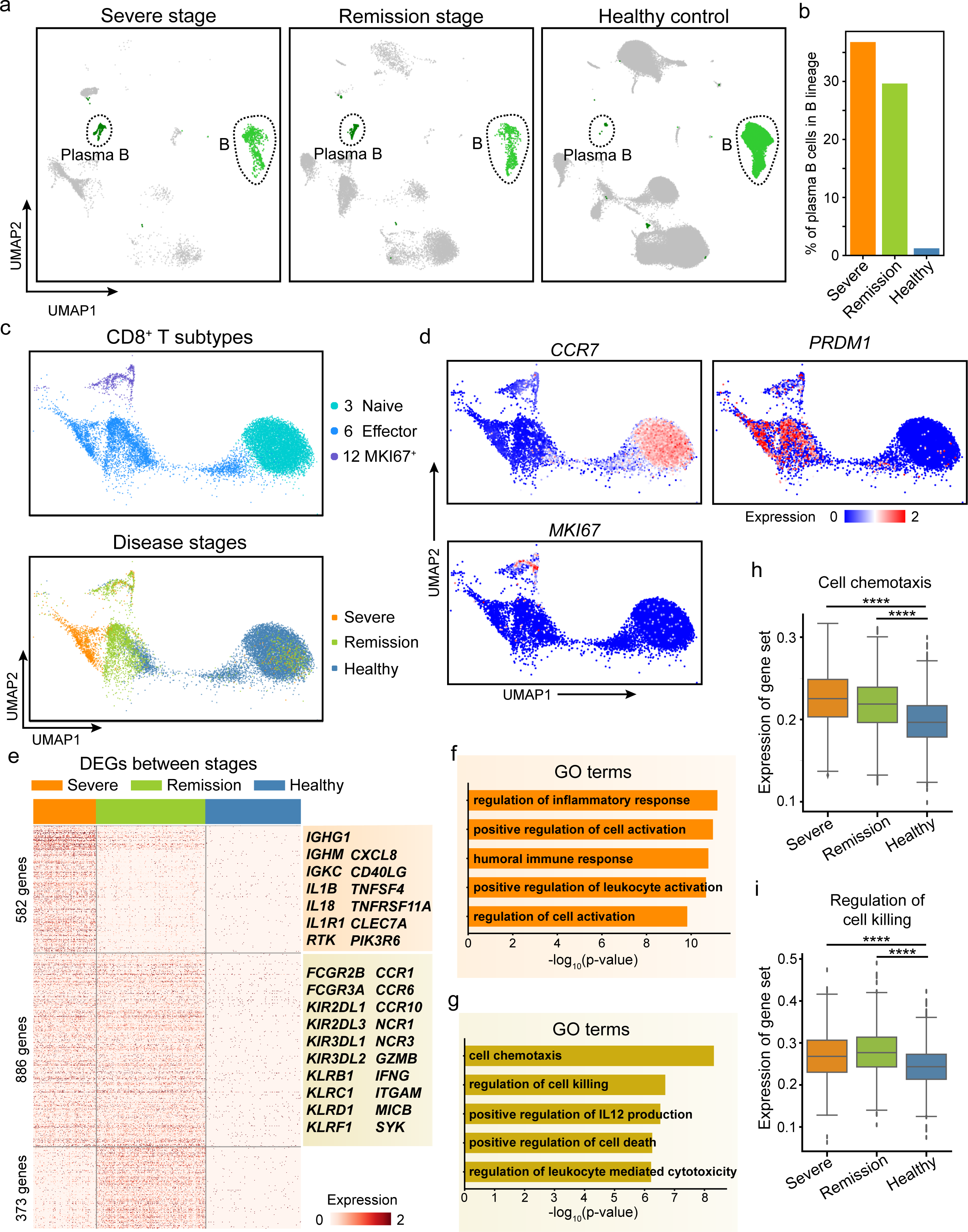
Enhanced humoral and cell-mediated immunity in severe COVID-19 patients. **a**, UMAP representations of B and plasma B cell clusters from the severe and remission stages, and in healthy control individuals. **b**, Bar plot of the proportions of plasma B cells in the B cell lineage from the severe and remission stages, and in healthy control individuals. **c**, UMAP representations of CD8^+^ T cell subtypes (left) and the distribution of cells from the severe and remission stages, and in healthy control individuals in each subtype (right). **d**, Dot plot of the expression of the *CCR7, PRDM1*, and *MKI67* genes in all CD8^+^ T cell subtypes. **e**, Heatmap of differentially expressed genes in effector CD8^+^ T cells from pairwise comparisons between the severe stage patients, remission stage patients, and healthy control individuals. **f, g**, Bar plots of GO terms enriched in effector CD8^+^ T cells from the severe stage **(f)** or the severe and remission stages **(g). h, i**, Box plots of the average expression of genes “cell chemotaxis” and “regulation of cell killing” in the effector CD8^+^ T cells from severe stage, remission stage, and healthy controls. Center line, median; box limits, upper and lower quartiles; whiskers, 1.5x interquartile range; points, outliers; **** represents *P* value < 10^−30^. Wilcoxon rank-sum test.

CD8^+^ T cells are function in cell-mediated immunity against viral infections by killing infected cells and secreting proinflammatory cytokines^30^. Our single-cell analysis detected a total of 13,602 CD8^+^ T cells. Clustering of these cells revealed 3 subtypes: naïve CD8^+^ T cells (cluster 3), effector CD8^+^ T cells (cluster 6), and a subset of CD8^+^ T cells with obvious expression of known proliferation markers (cluster 12) (Fig. 4c, d). The CD8^+^ T cells of the severe patients were primarily of the effector CD8^+^ T cell cluster (Fig. 4c, d). We then conducted pairwise comparisons to identify DEGs in the effector CD8^+^ T cells among the severe and remission stage patients and in healthy controls (Fig. 4e, Supplementary Table 8). A GO analysis indicated that DEGs in severe stage effector CD8^+^ T cells exhibited enrichment for “positive regulation of cell activation” (Fig. 4f, *P* < 10^−10^; hypergeometric test; Supplementary Table 9). Conversely, DEGs of the CD8^+^ T cells from severe and remission stage COVID-19 patients (i.e., vs. healthy controls) were enriched for functional annotations relating to “cell chemotaxis” and “regulation of cell killing” (Fig. 4g, *P* < 10^−6^; hypergeometric test; Supplementary Table 9). We also detected significant elevated expression of the 306 and 94 genes involved in these GO terms (Fig. 4h, i, *P* < 10^−32^, Wilcoxon rank-sum test; Supplementary Table 10). Together, these results indicate that SARS-CoV-2 infection elicits robust adaptive immune responses and suggest that Tocilizumab treatment further promotes such responses.

To gather additional empirical support from COVID-19 patients, we downloaded the bulk RNA-seq data of PBMCs from 3 severe COVID-19 patients and 3 healthy controls^31^, and applied AutoGeneS^32^ to deconvolute the composition of cell clusters based on the signature genes identified in our single-cell analysis. Our results indicated that there were significantly more severe-stage-specific monocytes (cluster 9), plasma B cells (cluster 11), and proliferating CD8^+^ T cells (cluster 12) in severe COVID-19 patients compared with healthy controls (Supplementary Fig. 8a-c, *P* < 0.05, Student’s t-test), findings consistent with our main conclusions.

## Discussion

The immune system is exerts essential functions in fighting off viral infections^33,34^ Recent studies have indicated that monocytes can exacerbate and even be a primary factor in the mortality of COVID-19 by contributing to inflammatory storms^4^. In the present study, we used single-cell mRNA sequencing and discovered a specific monocyte subpopulation that may lead to the inflammatory storm in severe-stage COVID-19 patients. By analyzing the monocyte-centric cytokine/receptor complements and predicting interaction networks, we uncovered a severe-stage-specific landscape of peripheral immune cell communication which may drive the inflammatory storm in COVID-19 patients. Our identification of this monocyte subpopulation and these cytokine-storm-related cytokine/receptor provides mechanistic insights about the immunopathogenesis of COVID-19 and suggests the potential of these cytokine/receptor molecules as candidate drug targets for treating the disease.

There have long been questions about whether treatment with the immunosuppressive agent Tocilizumab may affect the body’s antiviral responses^35,36^. Our single-cell profiles illustrated a sustained humoral and cell-mediated anti-virus immune response in severe and remission stage COVID-19 patients. For example, Tocilizumab treatment of severe-stage COVID-19 patients retained a high proportion of plasma B cells with antibody-secreting functions and we found that the cytotoxicity and cytokine production of effector CD8^+^ T cells remained stable upon Tocilizumab treatment.

Our work represents a collaborative clinical/basic effort that does provide an unprecedented empirical window for studying single-cell resolution profiles from severe COVID-19 patients. Deconvolution analysis of published bulk RNA-seq data^31^ from 3 additional severe COVID-19 patients and healthy controls helps support our conclusions on the enrichment of severe-stage-specific monocytes and plasma B cells in severe-stage COVID-19 patients. We further integrated additional single-cell datasets from sepsis patients and found the severe-stage-specific monocytes we observed are unique to severe COVID-19. Based on the incorporation of diverse additional data, our study and empirical data provide actionable insights that will help the multiple research communities who are still fighting against the virus, including clinical physicians, drug developers, and basic scientists.

## Methods

### Human samples

Peripheral blood samples were obtained from two severe COVID-19 patients. The patient severity was defined by the “Diagnosis and Treatment of COVID-19 (Trial Version 6)” which was released by The General Office of the National Health Commission and the Office of the National Administration of Traditional Chinese Medicine. Patient P1 was defined as a severe patient for his peripheral capillary oxygen saturation (SPO2) <93% without nasal catheter for oxygen. Patient P2 was defined as critical ill for respiratory failure, multiple organ dysfunction (MOD) and SPO2 <93 without nasal catheter for oxygen. Two peripheral blood samples were obtained for patient P1 on day 1 and day 5, and three peripheral blood samples were obtained for patient P2 on day 1, day 5 and day 7. For both patients, peripheral blood samples of day 1 were collected within 12 hours of Tocilizumab administration, when the patients were still at severe stage. Our decision to obtain blood draws from the two patients at day 5 were guided by information from the authors of the recent study published in *PNAS* ^13^, which guided our decision to consider day 5 of Tocilizumab treatment as a “remission stage”. For patient P2, we observed that his SARS-CoV-2 nucleic acid test of a throat swab specimen was still positive at day 5, so we took another blood draw at day 7 for P2, at point by which a throat swab specimen nucleic acid test was negative. All samples were collected from the First Affiliated Hospital of University of Science and Technology of China. Before blood draws, informed consent was obtained from each patient. Ethical approvals were obtained from the ethics committee of the First Affiliated Hospital of the University of Science and Technology of China (No. 2020-XG(H)-020).

### Cell Isolation

We collected 2ml peripheral blood each time from the COVID-19 patients. Peripheral blood mononuclear cells (PBMC) were freshly isolated from the whole blood by using a density gradient centrifugation using Ficoll-Paque and cryopreserved for subsequent generation of single-cell RNA library.

### Single-cell RNA-seq

We generated single-cell transcriptome library following the instructions of single-cell 3’ solution v2 reagent kit (10x Genomics). Briefly, after thawing, washing and counting cells, we loaded the cell suspensions onto a chromium single-cell chip along with partitioning oil, reverse transcription (RT) reagents, and a collection of gel beads that contain 3,500,000 unique 10X Barcodes. After generation of single-cell gel bead-in-emulsions (GEMs), RT was performed using a C1000 Touch^™^ Thermal Cycler (Bio-Rad). The amplified cDNA was purified with SPRIselect beads (Beckman Coulter). Single-cell libraries were then constructed following fragmentation, end repair, polyA-tailing, adaptor ligation, and size selection based on the manufacturer’s standard parameters. Each sequencing library was generated with unique sample index. Libraries were sequenced on the Illumina NovaSeq 6000 system.

### Single-cell RNA-seq data processing

The raw sequencing data of patients and health donors were processed using Cell Ranger (version 3.1.0) against the GRCh38 human reference genome with default parameters, and data from different patients and disease stages were combined by the Cell Ranger ‘aggr’ function. We are uploading the scRNA-seq data of PBMCs from the 2 severe COVID-19 patients to the Genome Sequence Archive at BIG Data Center and the accession number will be available upon request. We also used the scRNA-seq data of PBMCs from 2 healthy donors, which can be downloaded from the 10X genomics official website. Firstly, we filtered low quality cells using Seurat^15^ (version 3.1.4). For cells from COVID-19 patients (P1 and P2), we retained cells with detected gene numbers between 500 and 6,000 and mitochondrial unique molecular identifiers (UMIs) less than 10%. For cells from healthy donors, we retained cells with detected gene numbers between 300 and 5,000 and mitochondrial UMIs less than 10%. Subsequently we adopted Scrublet^14^ (version 0.2.1) to eliminate doublets in the PBMCs from the COVID-19 patients and healthy donors. We used the default parameters for Scrublet (i.e. “min_gene_variability_pctl=85, n_prin_comps=30, threshold=0.25”) and detected 50 doublets from the patients and 997 doublets from the healthy donors. After removing the doublets, we normalized gene counts for each cell using the “NormalizeData” function of Seurat with default parameters.

In downstream data processing, we used canonical correlation analysis and the top 40 canonical components to find the “anchor” cells between patients and healthy controls. We then used the “IntegrateData” function in Seurat to integrate cells from COVID-19 patients and healthy controls. We clustered all the cells based on the integrated gene expression matrix using Seurat with parameter “resolution=0.3” and generated 20 clusters. To display cells in a 2-dimensional space, we ran the principal component analysis on the integrated dataset and adopted the first 50 principal components (PCs) for the uniform manifold approximation and projection (UMAP) analysis.

In the integration of cells from COVID-19 and sepsis patients using Seurat, we applied the same functions and parameters as described above. We adopted Seurat to cluster the integrated gene expression matrix (with resolution = 0.3) and identified monocyte clusters based on the expression of known marker genes *CD14* and *CD68*. We then extracted all the monocytes from the integrated dataset and re-clustered them. Finally, we generated 10 cell clusters.

### Integration of cells from patients and healthy controls by Harmony

To verify the reliability of the integration results obtained using Seurat (version 3.1.4), we also applied Harmony^17^ to integrate the PBMCs from COVID-19 patients and healthy controls. We used the same gene expression matrix and applied the same parameters as Seurat, and adopted the first 50 PCs to perform the data integration by calling the “RunHarmony” function in Harmony. We then used the same clustering algorithm as Seurat to cluster cells and generated 23 clusters (“resolution=0.5”) based on the integration results obtained from Harmony. Jaccard index was applied to gauge the similarity between the cell clusters, with cell integration processed by Seurat or by Harmony. The Jaccard similarity between each pair of Seurat cluster (cluster *i*) and Harmony cluster (cluster *j*) were difined as follows

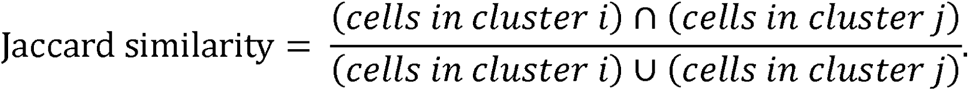

### Differential expression analysis

To search for the differentially expressed genes (DEGs), we first assign the negative elements in the integrated expression matrix to zero. We used Wilcoxon rank-sum test to search for the DEGs between each pair of the 3 stages of cells (i.e. severe stage, remission stage and healthy control). We applied multiple thresholds to screen for DEGs, including mean fold change >2, *P* value <0.001, and were detected in >10% of cells in at least one stage.

We defined stage A specific-DEGs as the intersections between the DEGs in stage A versus stage B and the DEGs in stage A versus stage C. We defined stage A and B common-DEGs as the intersections of the DEGs in stage A versus stage C and the DEGs in the stage B versus stage C, minus the DEGs between stage A and B. In this way, we obtained the specific-DEGs for each stage, and the common-DEGs for each pair of the 3 stages. We then uploaded these DEG groups to the Metascape^37^ website (https://metascape.org/gp/index.html#/main/step1), and used the default parameters to perform Gene Ontology (GO) analysis for each stage.

### Motif enrichment and regulatory network

We adopted SCENIC^19^ (version 1.1.2) and RcisTarget database to build the gene regulatory network of CD14^+^ monocytes. Since the number of CD14^+^ monocytes from healthy control (N = 9,618) was more than those from the severe and remission stages (N = 1,607), to balance their contributions in the motif analysis, we randomly sampled 2,000 CD14^+^ monocytes from the healthy control for calculation. We selected 13,344 genes that were detected in at least 100 monocytes or included in the DEGs of the 3 stages as the input features for SCENIC. With default parameters, SCENIC generated the enrichment scores of 427 motifs. We used the student’s t-test to calculate the *P* values of these motifs between severe stage and healthy control, and selected severe-specific enriched motifs with fold change >1.5 and *P* value < 10^−100^.

We then applied the enrichment scores of the severe-specific enriched motifs and the expression of their targeted genes to Cytoscape^38^ to construct a connection map for the gene regulatory network, as shown in Supplementary Fig. 6. The thickness of line connecting TFs and target genes represented the weight of regulatory link predicted by SCENIC.

### Cytokine/receptor interaction analysis

To identify potential cellular communications between monocytes and other cell types (CD4^+^ T, CD8^+^ T, B, plasma B and NK cells), we applied the CellphoneDB^26^ algorithm to the scRNA-seq profiles from the the severe and remission stages, and in healthy control individuals. CellphoneDB evaluated the impact of a ligand/receptor interactions based on the ligand expression in one cell type and its corresponding receptor expression in another cell type. We focused on the enriched cytokine/receptor interactions in severe-stage COVID-19 patients, and selected the cytokine/receptor interactions with more significant (*P* value < 0.05) cell-cell interaction pairs in the severe stage than that in the remission and healthy stages. We also included cytokine/receptor pairs which were highly expressed in severe stage.

### Deconvolution of cell clusters from bulk RNA-seq data

We applied AutoGeneS^32^ to deconvolute the composition of cell clusters based on the signature genes identified in our single-cell analysis. Specifically, we first obtained a gene-by-cluster expression matrix from our normalized single-cell profile, where the matrix elements were the average expression of each gene in each cell cluster. We then defined the top 5000 most variable genes between cell cluster and the 2000 DEGs used for cell clustering as the “VarGenes”, and extracted the VarGenes-by-cluster expression matrix as the feature gene expression profile for AutoGeneS. We set the input parameters as “model = ‘nusvr’, ngen = 1000, seed = 0, nfeatures = 1500” to deconvolute the cell composition in AutoGeneS.

### Statistical analysis

The two-tailed Wilcoxon rank-sum test (also called the Mann-Whitney U test) was used to search for the DEGs and to compare the expression differences of a gene set of interest between two conditions. In CellphoneDB, a permutation test was used to evaluate the significance of a cytokine/receptor pair. Metascape utilizes the hypergeometric test and Benjamini-Hochberg *P* value correction algorithm to identify the ontology terms that contain a statistically greater number of genes in common than expected. We used the Student’s t-test to evaluate the significance of the expression differences of the TFs (and their target genes) between samples from severe stage patients and healthy controls.

### Data Availability

The scRNA-seq data of PBMCs from the 2 severe COVID-19 patients can be obtained from the Genome Sequence Archive (GSA) at BIG Data Center and the accession number is CRA002509. We also used published datasets as controls or comparable data, including (1) the scRNA-seq data of PBMCs from 2 healthy donors downloaded from the 10X genomics official website https://support.10xgenomics.com/single-cell-gene-expression/datasets/3.1.0/5k_pbmc_NGSC3_aggr, (2) the scRNA-seq data of PBMCs from 22 sepsis patients and 19 related controls^25^ that is available on Institute Single Cell Portal (https://singlecell.broadinstitute.org/single_cell) with accesson number SCP548, (3) the bulk RNA-seq data of PBMCs from 3 COVID-19 patients and 3 related controls^31^ downloaded from the GSA at BIG Data Center with accession number CRA002390.

## Code Availability

Analysis scripts are accessible from github: https://github.com/QuKunLab/COVID-19.

## Funding

This work was supported by the National Key R&D Program of China (2017YFA0102900 to K.Q.), the National Natural Science Foundation of China grants (91940306, 81788101, 31970858, 31771428 and 91640113 to K.Q., 31700796 to C.G. and 81871479 to J.L.), the Fundamental Research Funds for the Central Universities (to K.Q.). We thank the USTC supercomputing center and the School of Life Science Bioinformatics Center for providing supercomputing resources for this project. We thank the CAS interdisciplinary innovation team for helpful discussion.

### Author Contributions

K.Q. conceived and supervised the project; K.Q., C.G. and J.L. designed the experiments; C.G. and J.L. performed the experiments and conducted all the sample preparation for next-generation sequencing with the help from H.M. and T.C.; B.L. performed the data analysis with the help from P.C., Q.Y., L.Z., L.J., C.J., Q.L., D.Z., W.Z., Y.L., K.L., X.G. and J.F.; T.C., X.W., L.L., J.W. and X.M. provided COVID-19 blood samples and clinical information; J.W. contributed to the revision of the manuscript; K.Q., C.G., J.L. and B.L. wrote the manuscript with the help of B.F., H.W. and all the other authors.

## Supplementary Figure Legends and Supplementary Tables

**Supplementary Figure 1.**
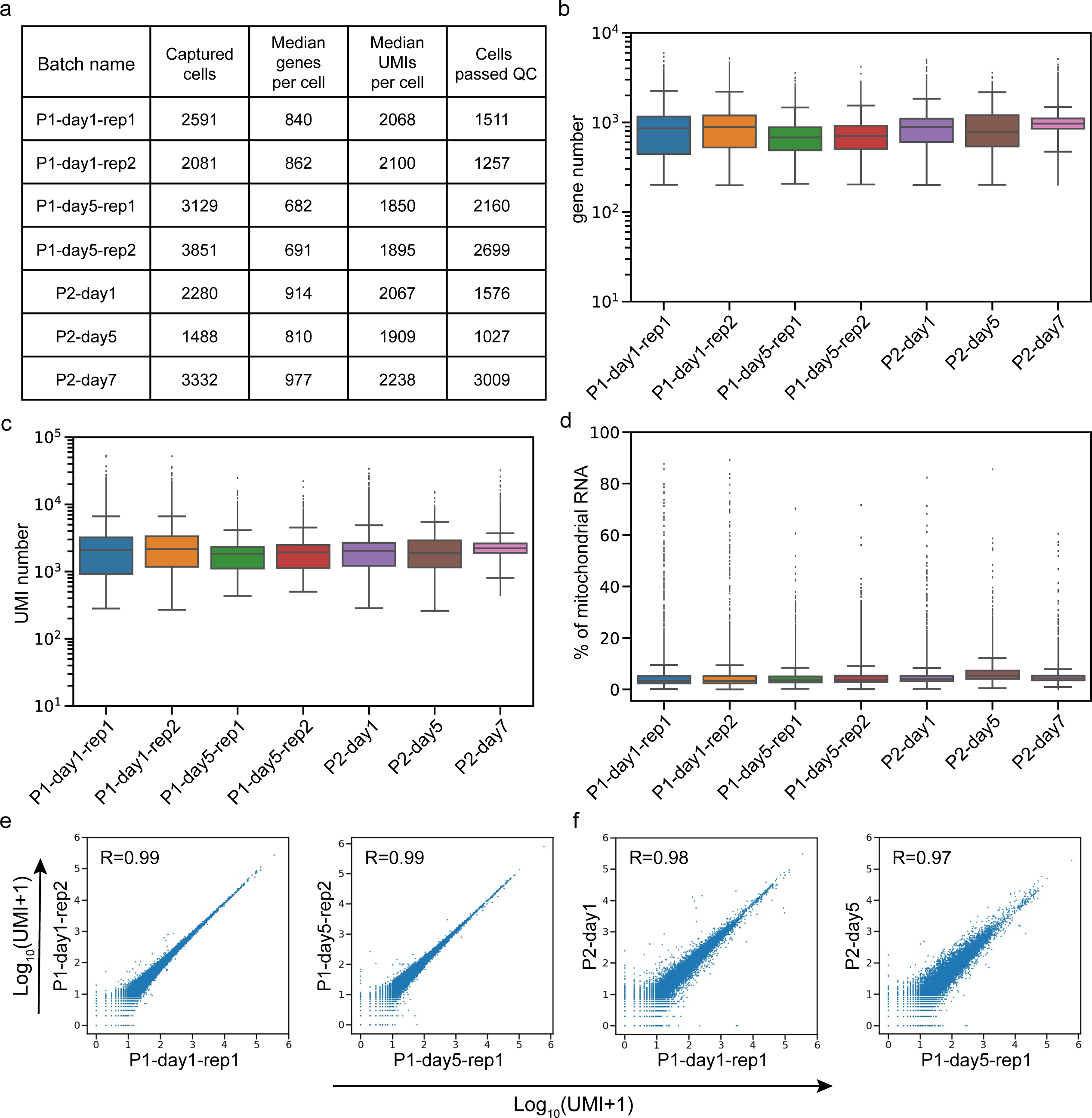
Quality control of single-cell data for PBMC samples from severe COVID-19 patients. **a**, Summary of captured cells, median genes per cell, median UMIs per cell, and the number of cells that passed quality control (QC) in distinct batches of single-cell data from severe COVID-19 patients. **b-d**, Box plots showing the gene number **(b)**, UMI number **(c)**, and percentage of mitochondrial RNA **(d)** in distinct batches of single-cell data from severe COVID-19 patients. **e, f**, Aggregated scRNA-seq one-to-one reproducibility plots for technical replicates **(e)** and biological replicates **(f)**. The correlation (R) represents the Pearson correlation across all genes. Box-whisker plot; the lower whisker is the lowest value greater than the 25% quantile minus 1.5 times the interquartile range (IQR), the lower hinge is the 25% quantile, the middle is the median, the upper hinge is the 75% quantile and the upper whisker is the largest value less than the 75% quantile plus 1.5 times the IQR.

**Supplementary Figure 2.**
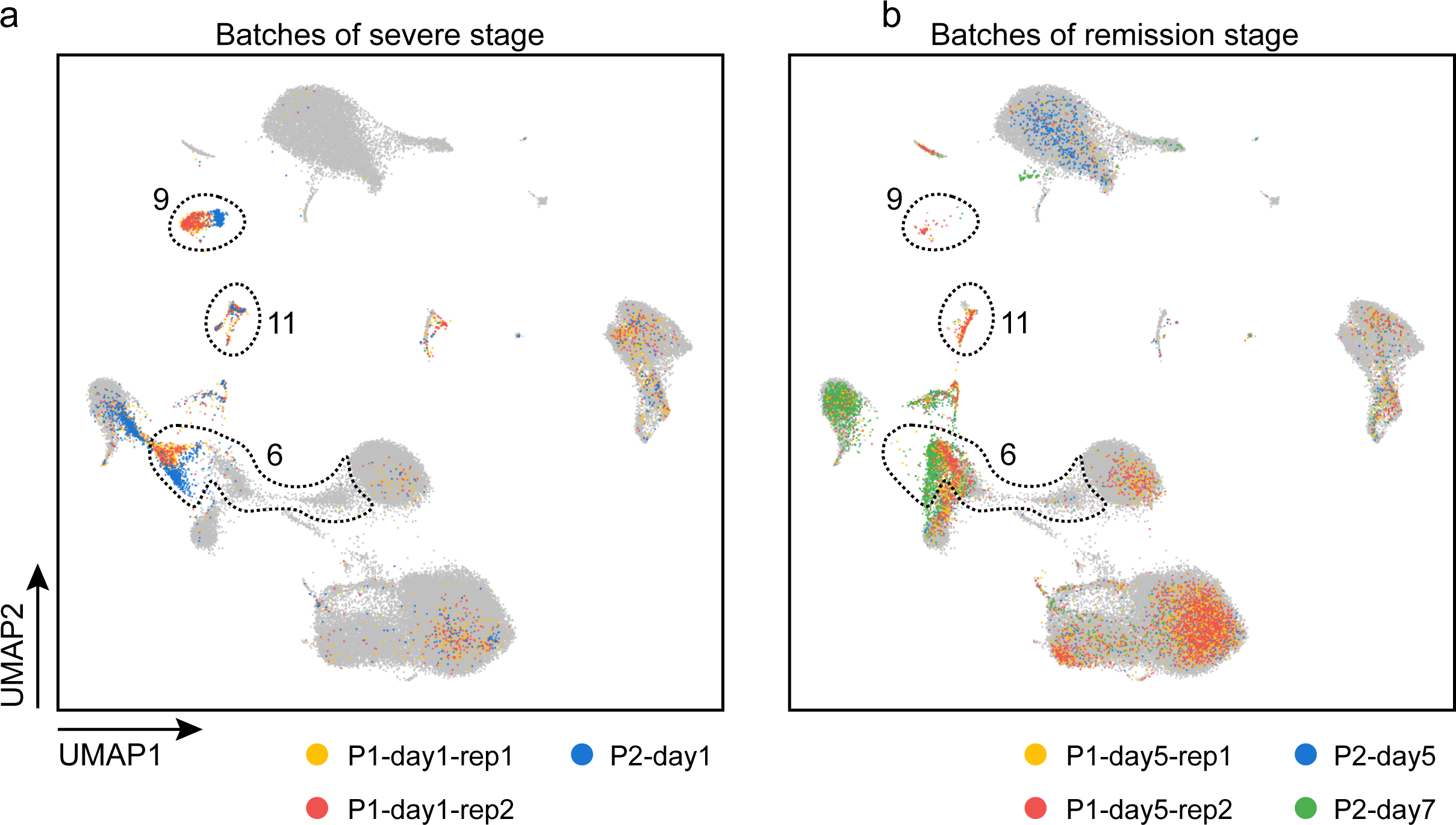
Single-cell transcriptomes of PBMCs from patient P1 or P2 at each time point. **a**, UMAP plot showing single-cell transcriptomes from patient P1 and P2 at day 1. **b**, UMAP plot showing single-cell transcriptomes from patient P1 at day 5 and P2 at day 5 and day 7.

**Supplementary Figure 3.**
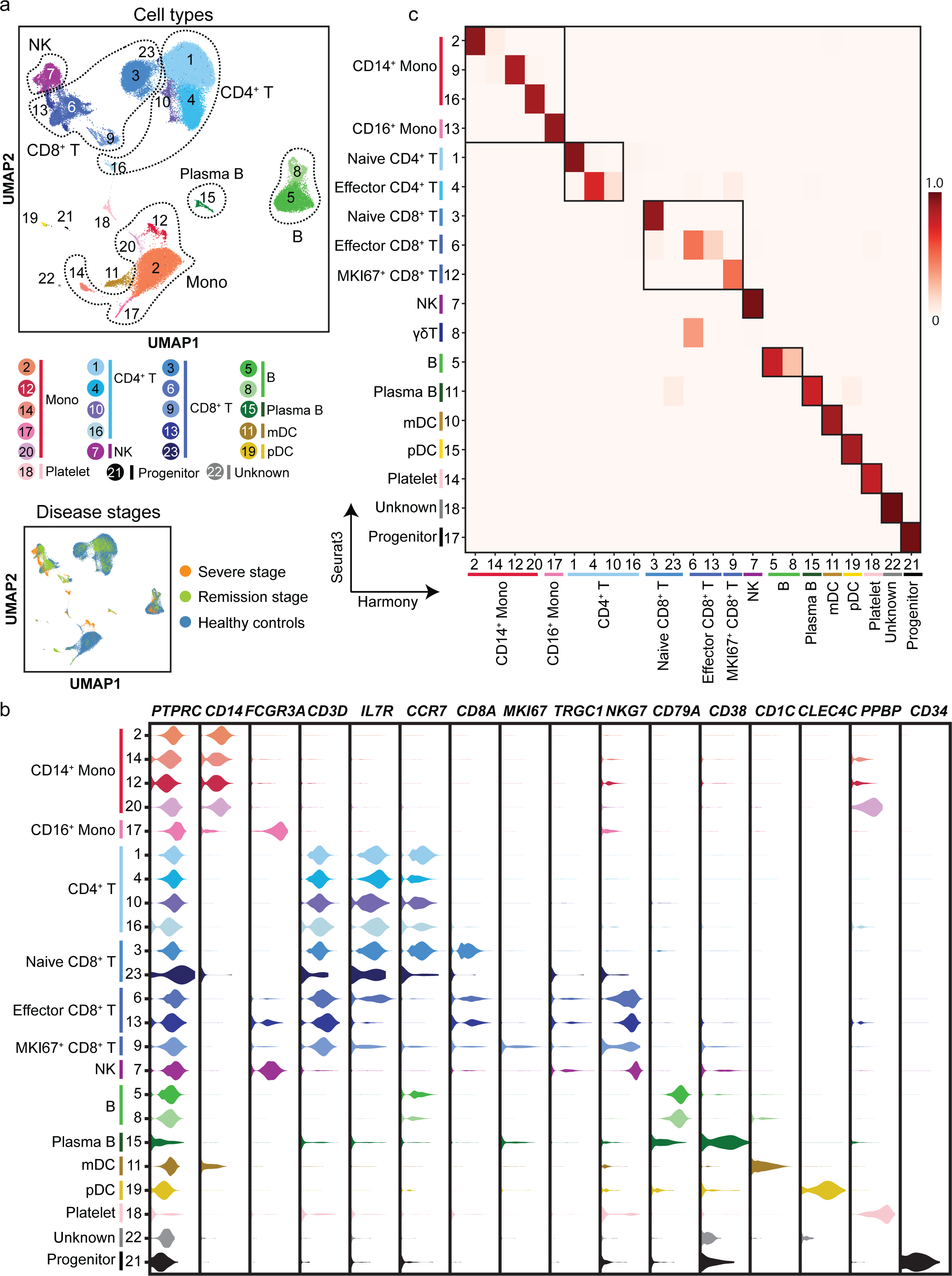
Single-cell profiling of peripheral immune cells in severe COVID-19 integrated with healthy controls using Harmony. **a**, UMAP representations of single-cell transcriptomes of 69,237 PBMCs integrated by Harmony. Cells are color-coded by clusters and disease state (see legend for key). Mono, monocyte; NK, natural killer cells; mDC, myeloid dendritic cells; pDC, plasmacytoid dendritic cells. **b**, Violin plots of selected marker genes (upper row) for multiple cell subpopulations. The left column presents the cell subtypes as identified based on combinations of marker genes. **c**, Jaccard similarities between the cell clusters with the integration processed by Seurat (version 3.1.4) and with the integration processed by Harmony.

**Supplementary Figure 4.**
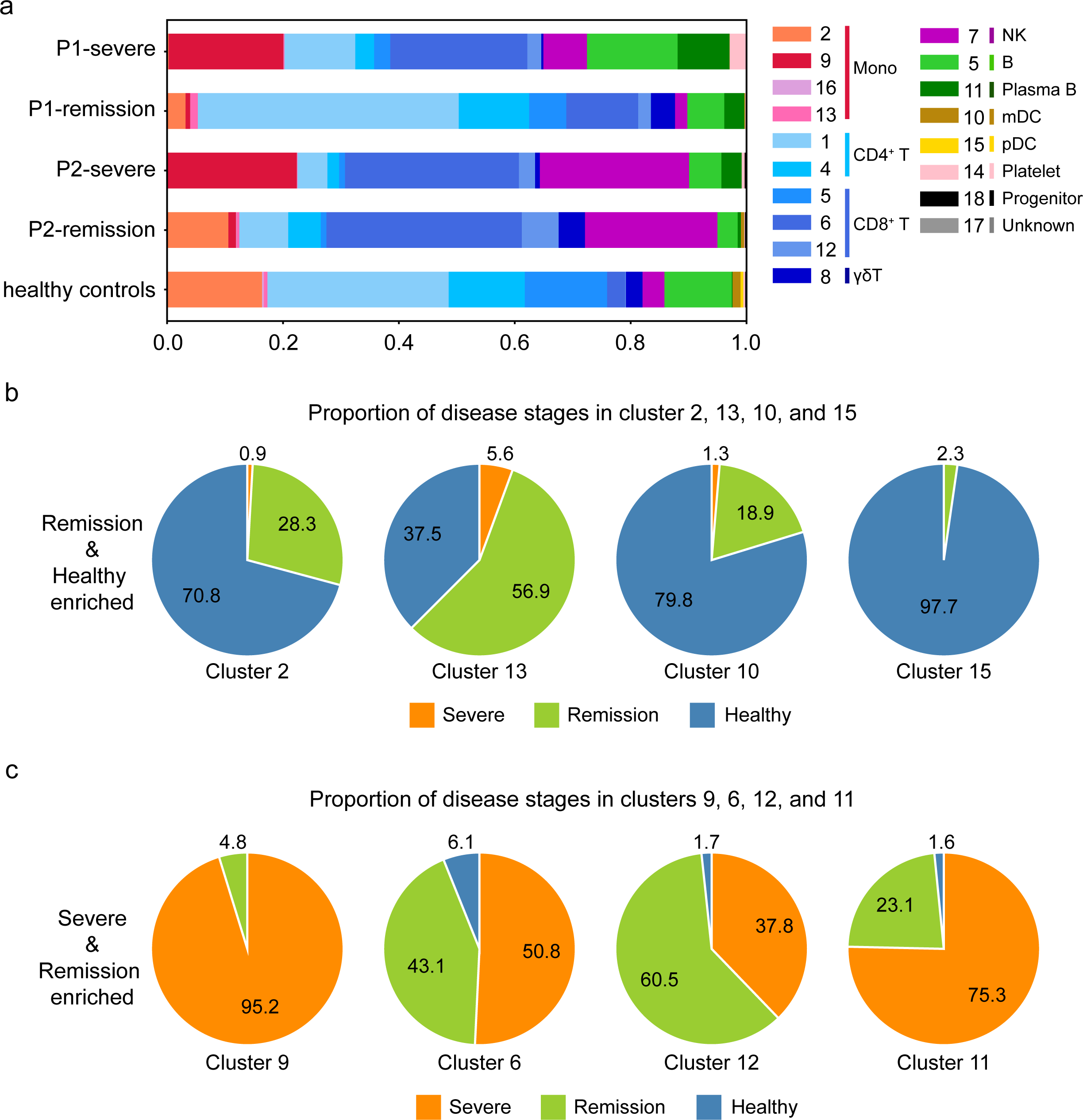
The composition of cell clusters identified in the integrated single-cell transcriptomes of PBMCs from the severe and remission stages, and in healthy control individuals. **a**, Bar chart showing the percentage of cell clusters in the severe and remission stages, and in healthy controls. **b**, Pie chart showing the proportion of cells from each disease state in selected cell clusters (cluster 2, 13, 10, 15), which were present in remission-stage patients and in healthy controls, but not in severe-stage patients. **c**, Pie chart showing the proportion of cells from each disease state in selected cell clusters (cluster 9, 6, 12, 11), which were present in severe and remission stages but not in healthy controls.

**Supplementary Figure 5.**
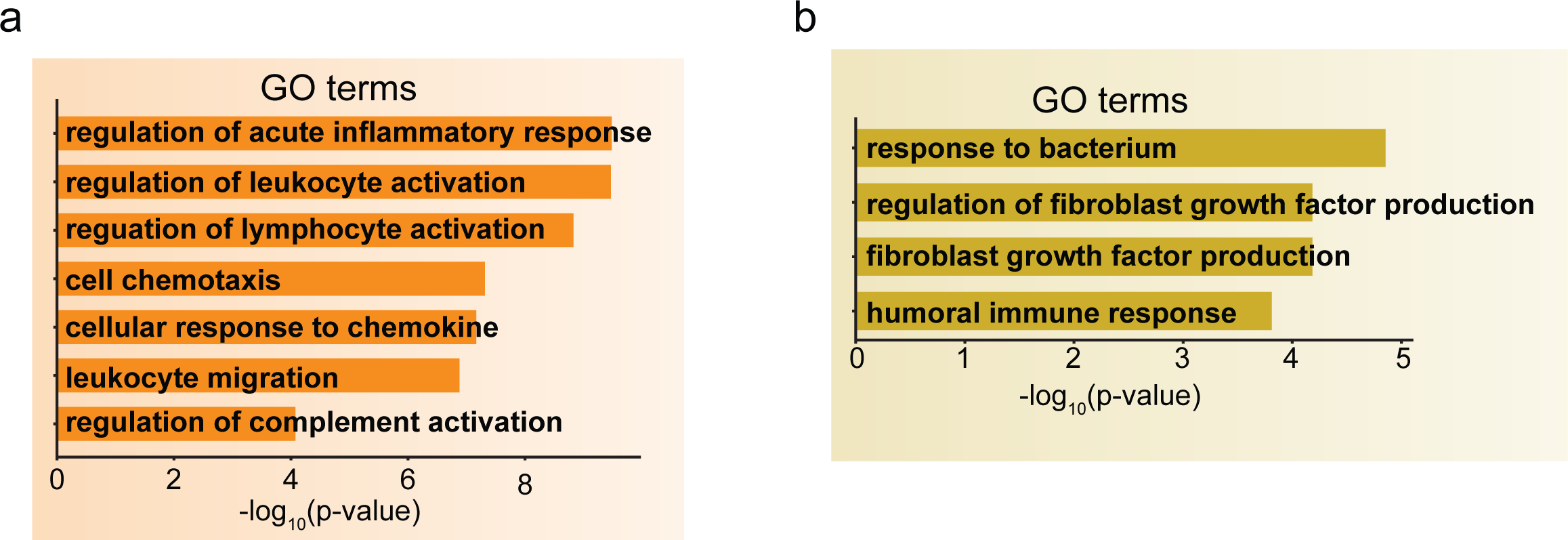
GO terms enriched among DEGs highly expressed in monocyte at the severe stage or at severe and remission stages. **a, b**, Bar plots of enriched GO terms of genes highly expressed in monocytes at the severe stage **(a)** or at the severe and remission stages **(b)**.

**Supplementary Figure 6.**
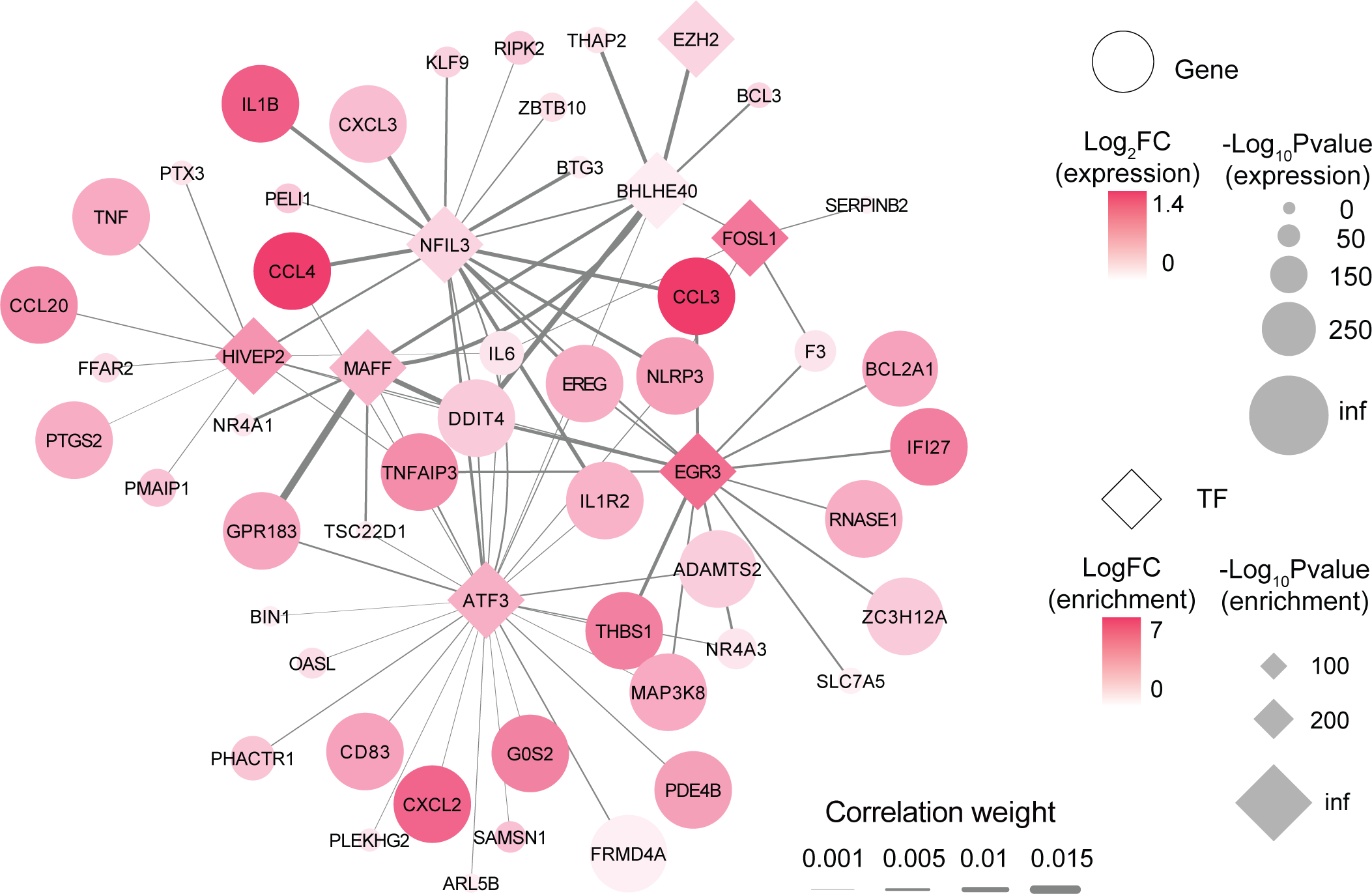
Severe-stage-specific monocyte regulatory network predicted by SCENIC. Transcription factors are shown as rectangles; their target genes are shown as circles. Student’s t-test.

**Supplementary Figure 7.**
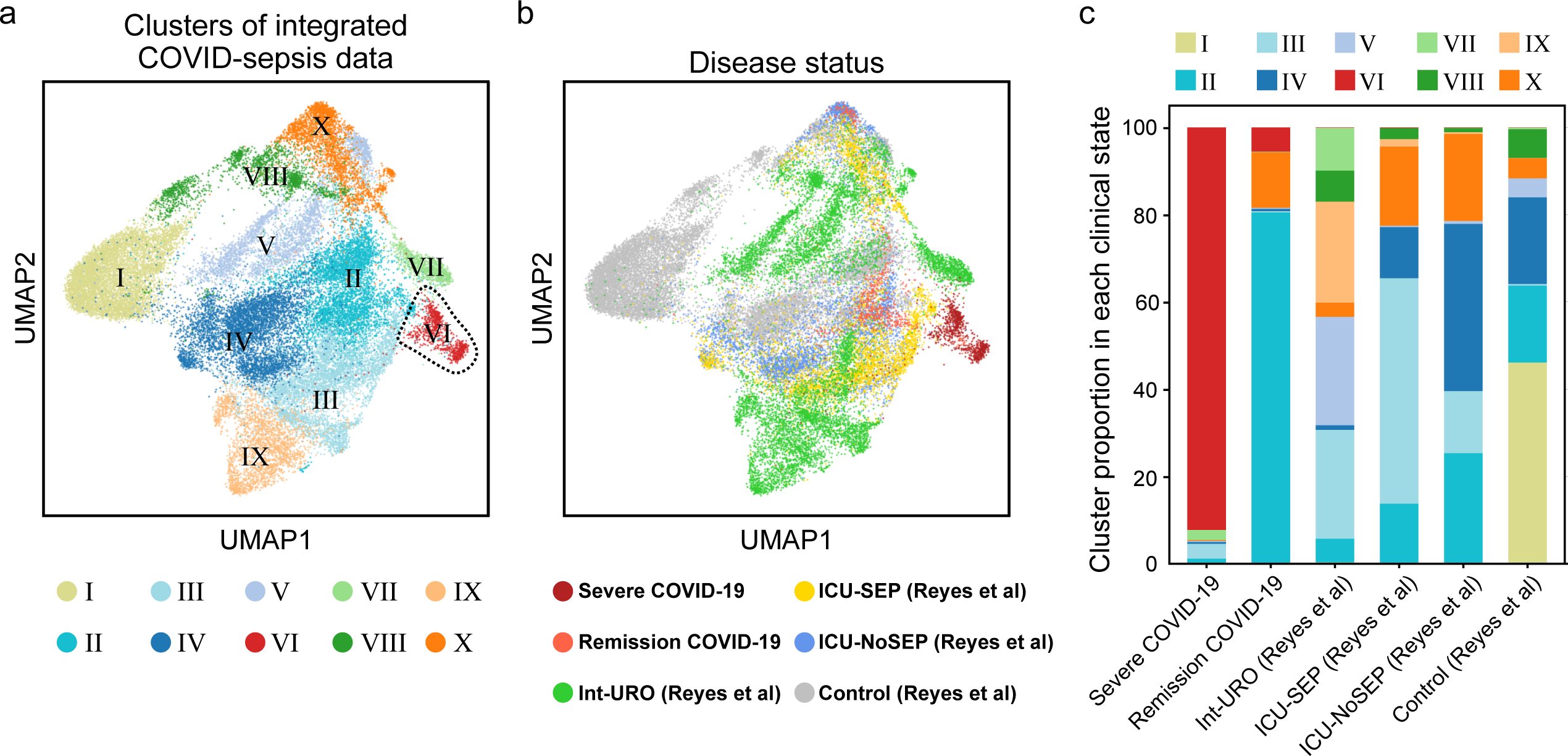
Integrated single-cell transcriptome analysis from patients with sepsis and our COVID-19 patients. **a, b**, UMAP representations of integrated single-cell transcriptomes from patients with sepsis at mild stage (Int-URO, n = 7)^25^, patients with sepsis at severe stage (ICU-SEP, n = 8)^25^, critically ill patients without sepsis (ICU-NoSEP, n = 7)^25^, healthy controls from outside our study (Control, n = 19)^25^, and our COVID-19 patients (Severe COVID-19 and remission COVID-19). Cells are color-coded by clusters **(a)**, disease states **(b). c**, Bar chart showing the proportion of cell clusters in (**a**) in each disease state.

**Supplementary Figure 8.**
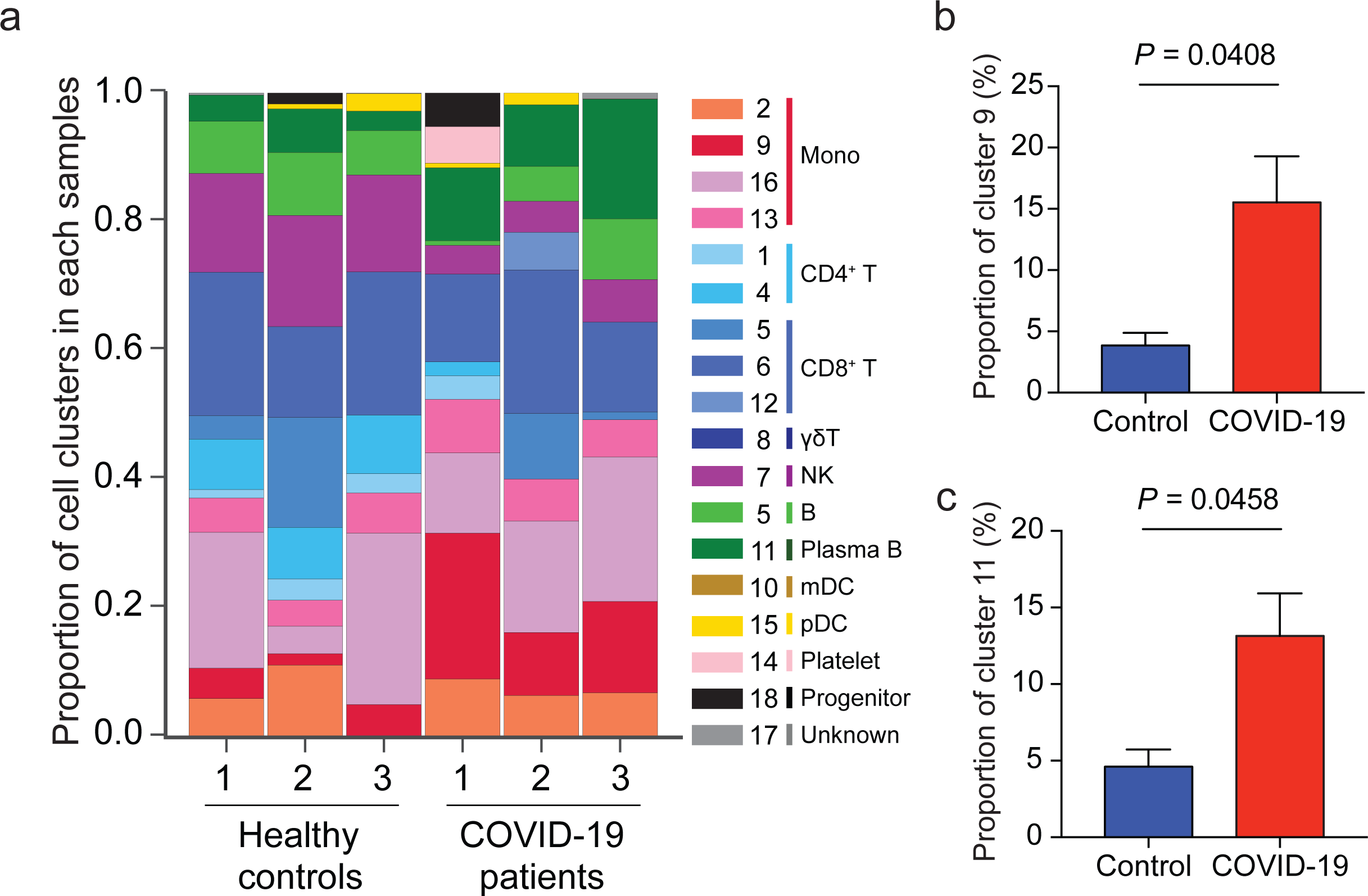
The composition of cell clusters identified in our single-cell analysis in a bulk RNA-seq from three severe COVID-19 patients and healthy controls. **a**, Bar chart showing an estimation of the composition of each cell cluster of PBMCs deconvoluted from bulk RNA-seq data from three COVID-19 patients and healthy controls^31^. **b, c**, Bar chart showing the percentage of severe-stage-specific monocytes (cluster 9, **b**) and plasma B cells (cluster 11, **c**) in COVID-19 patients and healthy controls, deconvoluted from bulk RNA-seq. Student’s t-test.

**Supplementary Table 1** | **Baseline characteristics and laboratory findings for the two COVID-19 patients in this study**.

**Supplementary Table 2** | **Sequencing data quality**.

**Supplementary Table 3** | **DEGs of different disease stages of the monocytes.**

**Supplementary Table 4** | **GO terms enriched among DEGs in different disease stages of the monocytes**.

**Supplementary Table 5** | **Sets of genes entailed in the enriched GO terms from Figure 2e and 2f**.

**Supplementary Table 6** | **Interactions of cytokines and receptors in different disease stages, predicted using CellphoneDB**.

**Supplementary Table 7** | **Drugs targeting cytokines or cytokine receptors.**

**Supplementary Table 8** | **DEGs of different disease stages of effector CD8^+^ T cells**.

**Supplementary Table 9** | **GO terms enriched among DEGs in different disease stages of the effector CD8^+^ T cells**.

**Supplementary Table 10** | **Set of genes entailed in the enriched GO terms from Figure 4h and 4i**.

