## Supplementary Table 1 for "Single-cell analysis of severe COVID-19 patients reveals a monocyte-driven inflammatory storm attenuated by Tocilizumab"

Baseline characteristcs and laboratory findings of COVID-19 patients in this study

| Patient ID | patient P1 | | patient P2 | | |
|---|---|---|---|---|---|
| Age, years | 39 | | 78 | | |
| Gender | male | | male | | |
| Chronic medical illness | steatohepatitis | | CHD, hypertension | | |
| date | Day1 | Day5 | Day1 | Day5 | Day7 |
| Disease stage | severe | remission | severe | remission | remission |
| Nucleic acid testing, (-) | positive | negative | positive | positive | negative |
| Cough, (-) | ++ | + | ++ | + | + |
| Expectoration, (-) | + | - | ++ | + | + |
| Chest distress, (-) | ++ | - | + | + | - |
| Chest CT grade | 15 | 14 | 15 | 14 | 11 |
| Peripheral capillary oxygen saturation, %, (95-100) (without nasal catheter for oxygen) | <93 | n/a | <93 | n/a | n/a |
| Nasal catheter for oxygen, L/min, (0) | 4 | 0 | 50 | 3 | 1- 2 |
| Peripheral capillary oxygen saturation, %, (95-100) (with nasal catheter for oxygen) | 97 | 98 | 95 | 100 | 99 |
| Heart rate, times/min | 72 | 82 | 86 | 115 | 85 |
| White blood cell count, X10^9/L, (4.00-10.00) | 5.03 | 4.91 | 8.90 | 6.20 | 4.46 |
| Lymphocyte percentage, %, (20-40) | 13.2 | 21.5 | 4.9 | 16.8 | 13.5 |
| Lymphocyte count, X10^9/L, (0.8-4.0) | 0.66 | 1.06 | 0.44 | 1.04 | 0.60 |
| Neutrophil percentage, %, (55-70) | 79.5 | 67.6 | 92.1 | 76.9 | 80.2 |
| Neutrophil count, X10^9/L, (2.5-7.5) | 4.00 | 3.31 | 8.20 | 4.77 | 3.58 |
| Haemoglobin, g/l, (120-165) | 147 | 138 | 139 | 126 | 112 |
| Platelet count, X10^9/L, (125-320) | 261 | 345 | 215 | 160 | 122 |
| C-reactive protein, mg/L, (0.8-8) | 48.30 | 0.50 | 25.20 | 1.5 | 0.5 |
| IL-6, pg/mL, (0-7) | 8.98 | 43.21 | 45.68 | 197.70 | / |
| Alanine aminotransferase, IU/L, (0-40) | 116 | 133 | 101 | 177 | 141 |
| Aspartate aminotransferase, IU/L, (0-40) | 56 | 51 | 48 | 109 | 82 |
| Total bilirubin, umol/L, (3.4-17.1) | 41.2 | 16.8 | 26.8 | 30.9 | 30.8 |

CHD, coronary heart disease
() Values in parentheses are referenced normal values
++ The symptom is severe
+ The symptom is mild
- The symptom is disapeared
/ The patient did not do this test on that day
